# ProteoMill: Efficient network-based functional analysis portal for proteomics data

**DOI:** 10.1101/2020.11.09.374579

**Authors:** M Rydén, M Englund, N Ali

## Abstract

Functional analysis has become a common approach to incorporate biological knowledge into the analysis of omics data, and to explore molecular events that govern a disease state. It is though only one step in a wider analytical pipeline that typically requires use of multiple individual analysis software. There is currently a need for a well-integrated omics analysis tool that performs all the steps. The ProteoMill portal is developed as an R Shiny application and integrates all necessary steps from data-upload, converting identifiers, to quality control, differential expression and network-based functional analysis into a single fast, interactive easy to use workflow. Further, it maintains annotation data sources up to date, overcoming a common problem with use of outdated information, and seamlessly integrates multiple R-packages for an improved user-experience. The functionality provided in this software can benefit researchers by facilitating the exploratory analysis of proteomics data.

ProteoMill is available for free at https://proteomill.com.

## Background

The large amounts of data generated from omics experiments have stressed the need for methods to reveal and extract critical components of dynamic biological systems in a readable manner, which connects to the specific study question. Expression data that are derived from high throughput analysis have multiple levels of biological features connected to it. In a real biological environment, the physical, genetic, regulatory, and functional properties of a molecular set work together in a response to environmental stimuli. Holistically evaluating these attributes is a way to reveal the intercommunication between these properties and to provide a biological context. However, this task encompasses some impending challenges, including differences in biomolecule identification, data dimensionality reduction, biological contextualization, statistical analysis and data visualization, and this differs among the various types of individual datasets.

The existing omics analysis tools are typically specialized for individual parts of the analysis workflow and are not integrated between each other. This requires the researcher not only to have knowledge of the different individual software, but also knowing how to format the generated output from one software for use in the next software. This often poses a time-consuming task, particularly for researchers with little computational experience, or little experience with the software(s) in question and is prone to errors. Another important but often overlooked aspect for generating reliable and biologically relevant results is the quality of annotation data, and, by extension, a tool’s ability to maintain annotation data sources up to date. Lina Wadi et. al reported that 67% of publications in their survey referenced software using outdated annotation data^1^. Web-based tools have an inherit advantage in that back-end data sources can be dynamically updated without requiring manual action by the user (such as downloading and installing software). While there are many existing web-based omics tools which are able to perform individual parts of an analysis workflow ^2–7^ there is still a need for fast, interactive web-tools that can perform the complete workflow.

Analysis of proteomic data faces additional challenges^8^. Different gene- and protein level identifier types are utilized in the various omics tools, which often require the researcher to convert between identifier types before proceeding to the next step of the analysis. This can result in loss of data since there can exist one-to-many mappings between two identifier types, or that an identifier cannot be mapped between two identifier types ^9^. Furthermore, a frequent concern in mass spectrometry-derived data is the abundance of missing values ^10, 11^.

Thus, a tool that could help to transform the biological research into integrated framework is preferred. The aim of this study is to describe a newly developed software that addresses many of the existing shortcomings. The fundamental concepts of this software are to provide sets of well-integrated, easy-to-use, and to a large extent automated functions for exploratory analysis of proteomic data.

## Results

### Core functionality and architecture

The overall architecture and dataflow in ProteoMill can be visualized (Fig. 1) as three separate modules: ***Local***, the storage on the user’s computer. Any files that are uploaded or downloaded reside here. ***Core***, the main functionalities that ProteoMill offers. This includes any function carried out on the server, such as differential expression analysis, or exporting a report. ***External***, any data sources that are hosted some place other than the user’s computer or the ProteoMill server, for example, the protein interaction data which is collected and processed from STRINGdb^12^.

**Figure 1.**
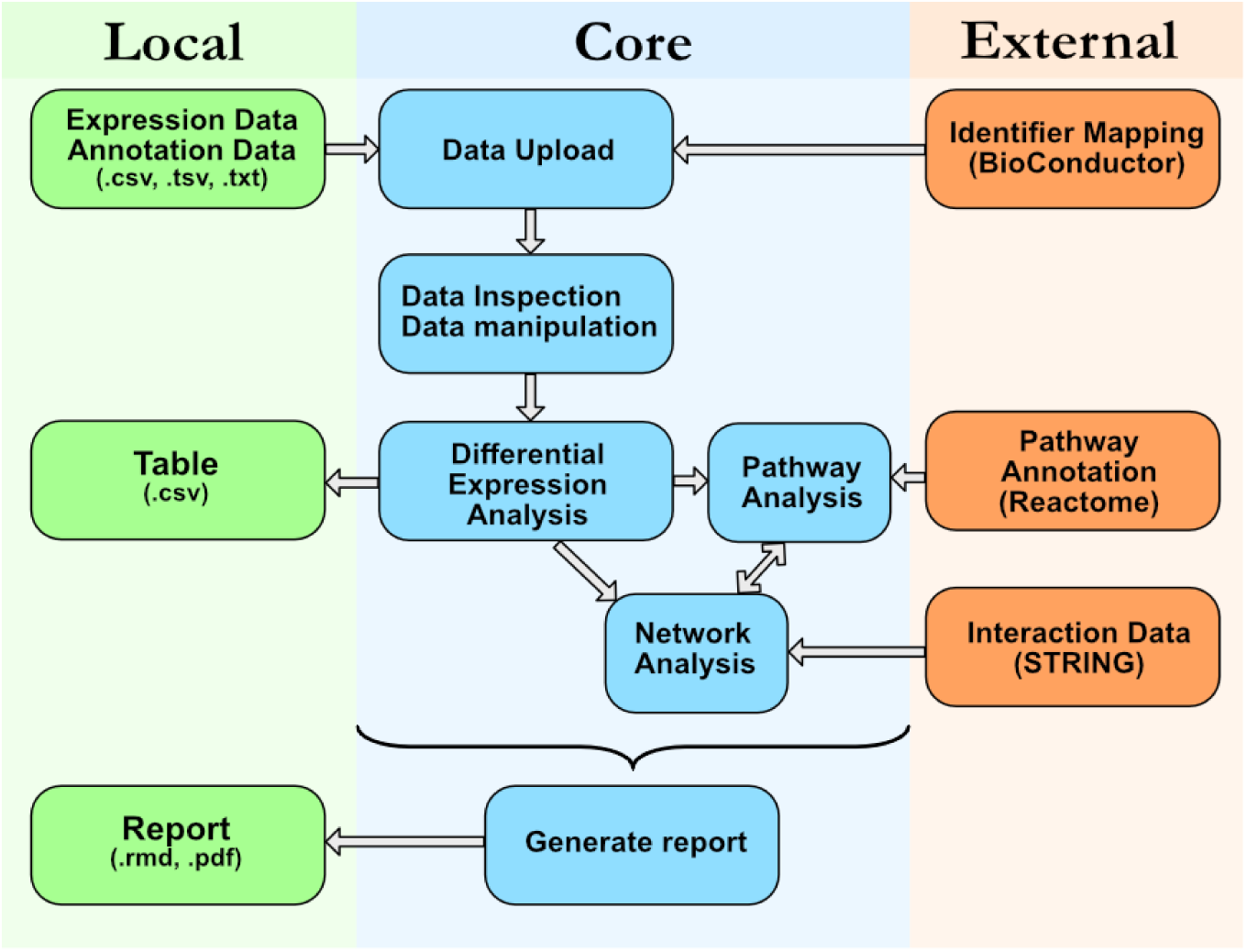
Schematic overview of core ProteoMill components and their relation to local and external environments.

### User interface

The user interface consists of a left-hand menu through which the user navigates through the analysis workflow. Adhering to the theme of automation, the data-upload section lets the user upload the main dataset and optional annotation data with a single click, with the optional settings column separator and identifier type. The user can also choose to use demo datasets.

Throughout the analysis, a task menu (Fig. 2) is dynamically updated with feature documentation and hints to progress. For instance, when the “Upload a dataset” task item is clicked, a guide displaying the expected data format is shown. This page contains a typical example, as well as a feature to design an example dataset based on the number of conditions, replicates and identifier type of the user’s choice. The task feature offers both easily accessible documentation, as well as an informative description of common methods such as differential expression.

**Figure 2.**
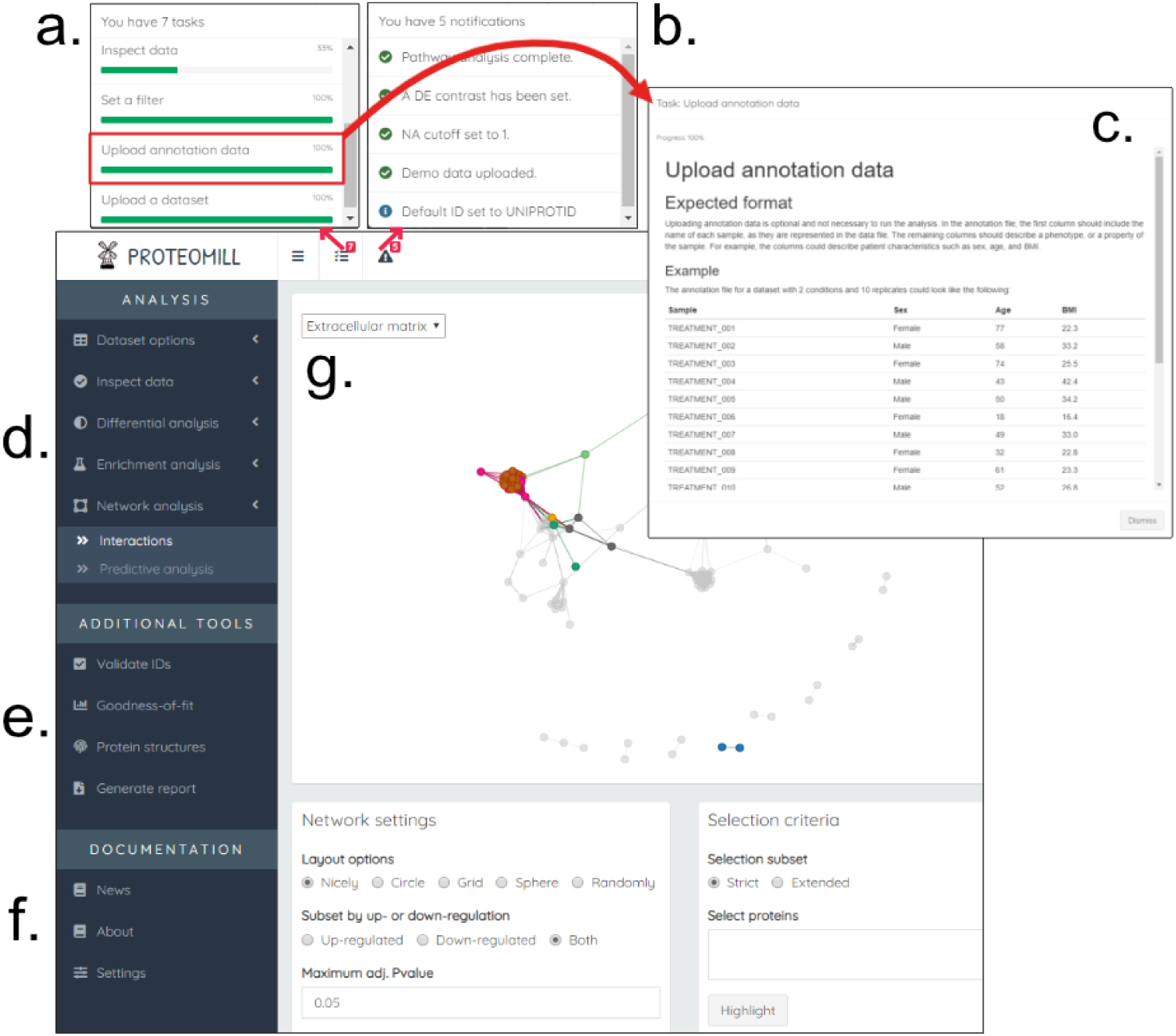
Task list. **a**. Each item describes the progress of the analysis workflow. **b**. Upon clicking an item, a window (*c*) is displayed with a thorough description of the task together with conceptual background of the analysis step. **d**. The analysis workflow menu, where each tab-item corresponds to a step in the workflow, and each sub-item corresponds to a particular function within that analysis step. **e**. Useful features which are not included in the standard analysis workflow. **f**. The documentation section includes news about ProteoMill, general information about the tool, and a settings menu. **g**. The main analysis area is where user-interactions and plot/table output are visible.

**Figure 3.**
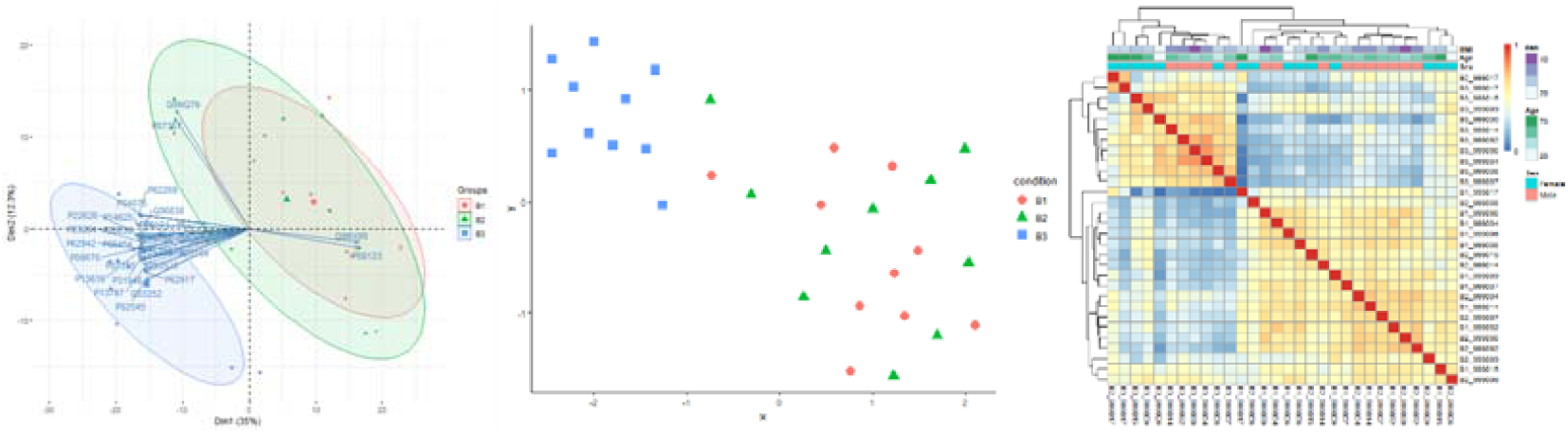
Visualization strategies for data inspection. **a**. PCA-biplot, which combines the information given by PCA and plot of loadings. **b**. UMAP is a fast, nonlinear dimensionality reduction method. **c**. Annotated sample-sample correlation heatmap.

The commitment to reproducibility and documentation is also manifested by a feature to generate a report, which can be done at any point during the analysis, to export the produced plots and tables as a markdown-formatted HTML, PDF, or Word-file.

### Data inspection and preprocessing

As missing values are a common obstacle in mass spectrometry-derived proteomic data analysis, ProteoMill incorporates a feature to visualize the number proteins affected by missing values (Fig. 8). The user can set a threshold for the maximum allowed missing values for each condition and exclude those proteins with too few quantitative values to be considered informative.

The UniProtKB identifier type is commonly used in proteomics data, but its identifiers can become obsolete (Fig. 4) unless the database is continuously updated ^13^. ProteoMill has a feature that lets the user validate the identifiers by displaying the identifier history, for detecting obsolete identifiers, which then can be updated. Through the Settings menu, the user can select between five identifier types to be used as the labels in plot- and table outputs and is dynamically updated upon selection.

**Figure 4.**
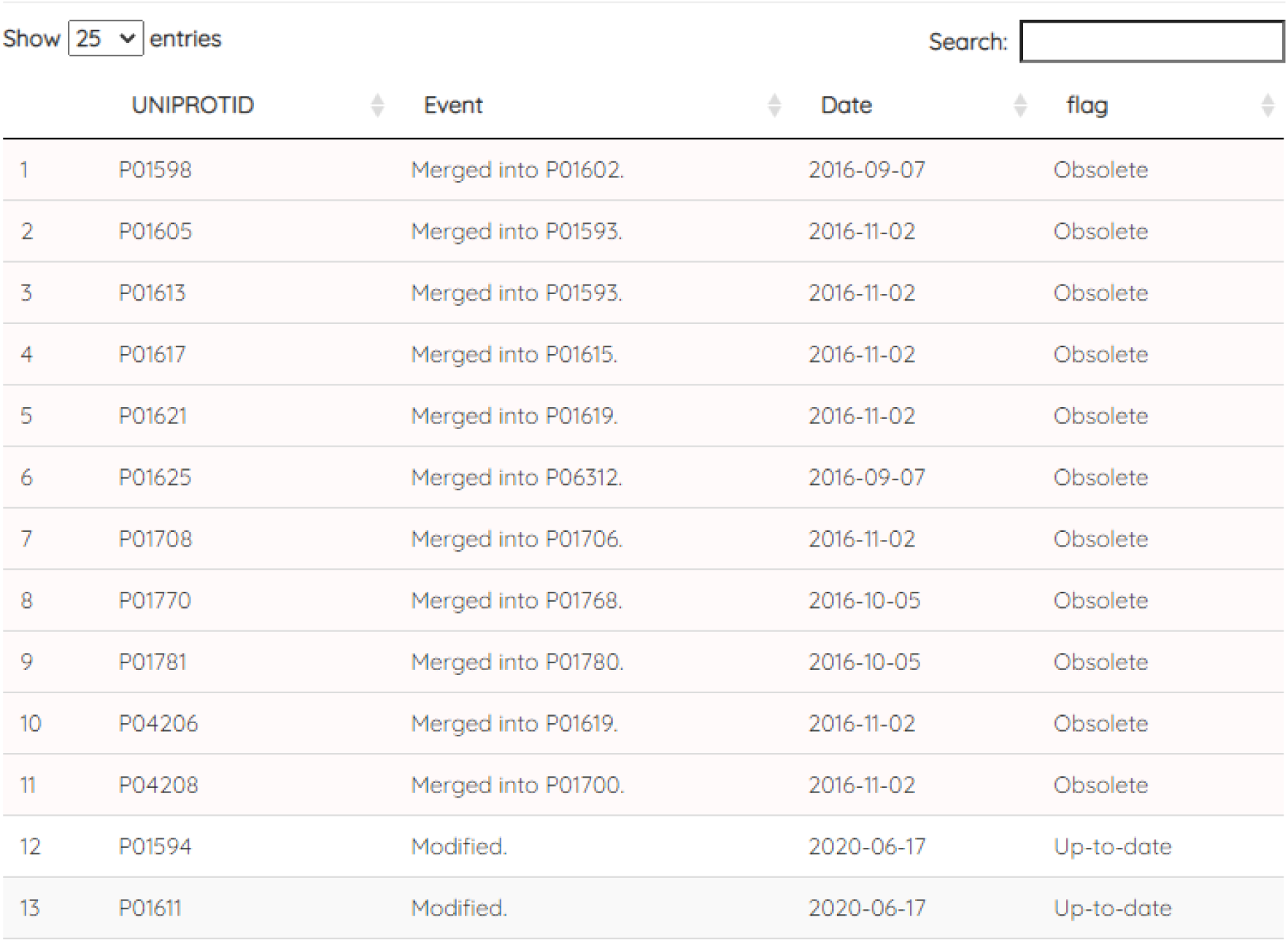
ProteoMill can retrieve the event history of a UNIPROTKB identifier. Here, it is used to determine if an identifier is obsolete or up to date.

The data inspection sub-menu consists the dimension reduction methods Principal Component Analysis (PCA) and Uniform Manifold Approximation and Projection (UMAP). PCA plots are implemented both as a 2D PCA-biplot, which displays the N proteins that contribute the most to the sample clustering effect, and a 3D PCA-plot, which can be inspected interactively. A sample-sample correlation heatmap can be generated to display sample clustering and quality-check the data. Additionally, if annotation data has been uploaded, this too will be displayed in the heatmap. In the case of the uploaded example data, three variables annotate the heatmap: BMI, age, and sex.

### Differential expression analysis

The packages limma^14^ and DESeq2^15^ are implemented for differential expression analysis. The researcher may first assess goodness-of-fit by visual inspection of quantile and density function plots generated by fitdistrplus^16^, to see which model is more appropriate for the uploaded dataset.

Differential expression analysis is conducted by specifying which two conditions should be contrasted, with an optional setting to specify a paired design. The output is displayed as a table with fold-changes, confidence intervals, and p-values for each protein from the differential expression results. (Table 1)

**Table 1.**
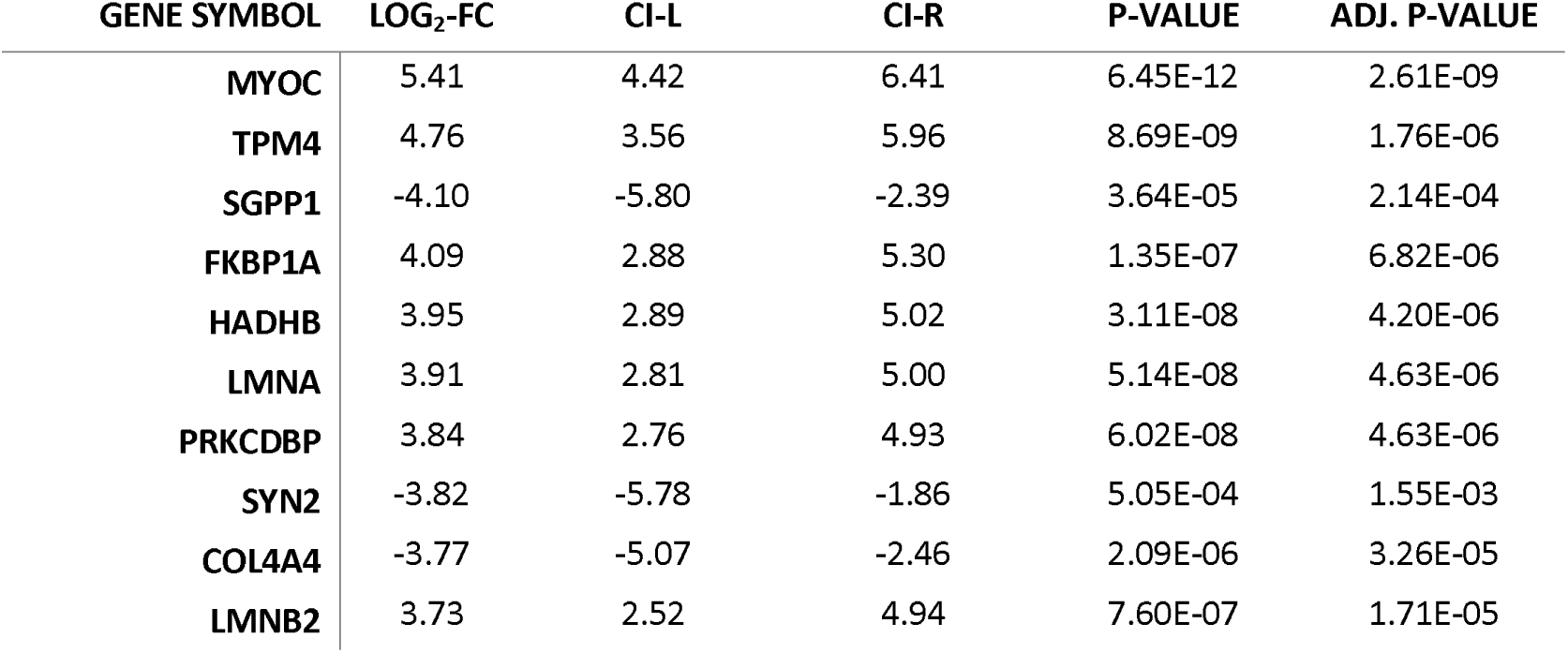
The ten proteins with highest absolute base-2 log fold change in the two conditions contrasted. The table was produced in ProteoMill.

### Functional analysis

The sets of up- and downregulated proteins generated from differential expression are used to explore which pathways the proteins are over-represented in. The results are displayed as two separate tables for up- and down-regulation, both containing the over-represented pathway name, its corresponding top-level pathway in the hierarchy, and a description of the relevance of the pathway.

This pathway over-representation results can be further visualized as a volcano plot (Fig. 5b), where each point is colored according to the most over-represented pathway it is annotated to.

**Figure 5.**
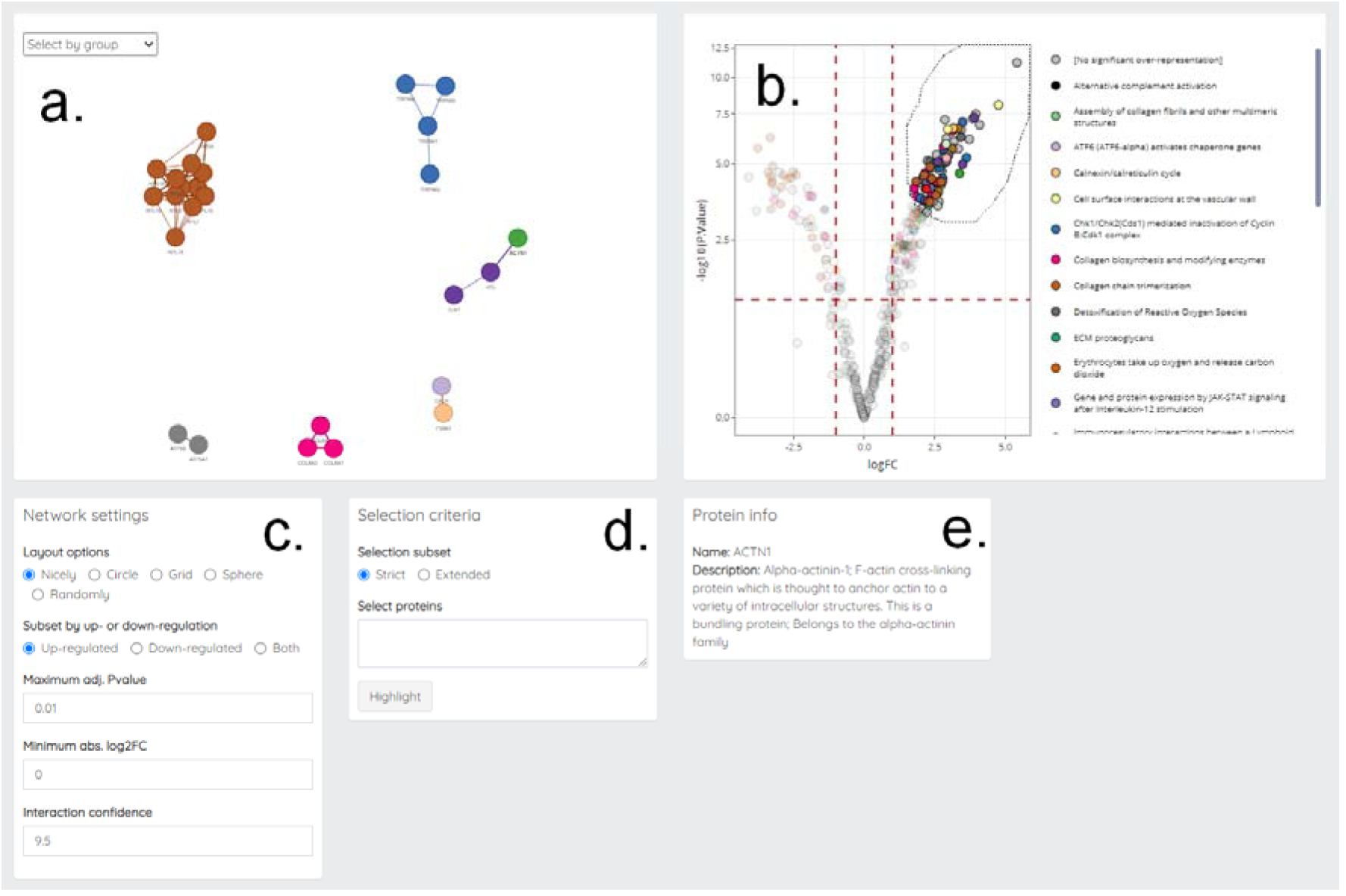
Visualization of functional enrichment results. **a**. Network of protein interactions, restricted according to parameters set in *b, c*, and *d*. **b**. Volcano plot. **c**. Network settings. **d**. Selection subset. “Strict” displays only interactions between nodes selected in *b*, while “Extended” displays interactions between nodes selected in *b* and any other protein in the current dataset. Nodes in *a* will be highlighted when their corresponding protein identifier is present in the “Select proteins” field. **e**. Protein info. Hovering over a node in *a* will display a summary of the protein’s function.

The network analysis tab has a feature that integrates the interactive volcano plot, so that users can select proteins of interest to see how the specific proteins interact. There exist multiple options to reshape the network to only include nodes of interest (Fig. 5c):

- Display only up-regulated or only down-regulated proteins.
- Subset the network based on differential expression significance level (include proteins lower than specified adjusted p-value).
- Subset the network based on differential expression fold-change (include proteins which have an absolute base-2 log fold change higher than the specified number).
- Show only protein-protein interaction that higher than set “confidence score” (as defined by Szklarczyk, D. *et al*., 2010).^12^

The network can further be reduced by using the selection tool in the interactive volcano plot (Fig. 5b) to circle-select nodes of interest. Any individual pathway listed next to the plot can be double-clicked to display only proteins annotated to that pathway. Moreover, under “Selection criteria” (Fig. 5d), one can select “Strict” to display only interactions between selected nodes, or “Extended” to display interactions between selected nodes and any other protein in the current dataset. A list of protein identifiers can be put into the “Select proteins” text area to highlight the nodes of corresponding proteins in the network area. Hovering over a node will display a summary of the protein’s function (Fig. 5e).

### Case study

We demonstrated the software’s capabilities on a previously published dataset^17^ that contained 638 proteins, identified and quantified by mass spectrometry based proteomics on human meniscus samples, from which a slice of the mid-portion of the meniscus body had been divided into three zones: inner (B1), middle (B2) and peripheral (B3). This dataset was used when producing all plots and tables in this paper.

Out of 638 initial proteins in the original dataset, 12 proteins had obsolete identifiers which were substituted to their updated equivalents. 405 proteins remained after setting a missing value threshold of one missing value per zone. Data inspection revealed that the B3 samples clustered together, separate from B1 and B2.

In the comparison of meniscal zones B3 and B1, 129 proteins had an absolute base-2 log fold-change greater than 2. Highest absolute base-2 log fold change was seen in myocilin (5.41), tropomyosin alpha-4 chain (4.76), trifunctional enzyme subunit beta (3.95), prelamin-A/C (3.91) and caveolae-associated protein 3 (3.84).

Enrichment of up-regulated proteins showed an over-representation of proteins in pathway categories related to metabolism of RNA and metabolism of proteins. For the down-regulated proteins, many over-represented pathways were related to extracellular matrix organization.

The results are in line with those described by Folkesson et. al in an article describing the same dataset.^17^

## Discussion

The presented software, ProteoMill, proposes a unique approach to conducting explorative analysis of proteomics data. Researchers who want to use this tool do not need programming experience and the platform is readily available through the browser.

The data visualization capabilities present in this software are designed to make it possible even for researchers without any particular computational training to gain insights about the biological meaning of their data. Many of the graphical components are interactive, which is a useful feature for analyzing protein interactions, and selecting subnetworks of interest.

A common goal in many of ProteoMill’s functionalities is to reduce data complexity, and to provide a framework for extracting elements of biological relevance. PCA reduces a dataset of hundreds or thousands of expression datapoints into a single datapoint for each condition, plotted in 2-3 principle components, which in turn describes the dimensions with largest variability. The datapoints cluster together based on the similarity of their expression profiles.

Categorizing proteins into biological entities, described as pathways, is another way to reduce complexity and make sense of one’s data. Network graphs produced from interaction data can be difficult to interpret. In ProteoMill, pathways are used to categorize and label groups of interacting proteins, and as a way to inspect subnetworks based on these common biological themes.

The integrated enrichment- and network analysis provides a way for users to simultaneously explore functional analysis output and interaction data, and this feature has been specifically designed to easily identify and select subnetworks of interest for further analysis.

A central aspect when developing the platform was the importance of regular updates of data sources. As discussed in a paper by Zhou et al ^18^, many platforms rely on databases which are out-of-date and have great impact on the outcome of enrichment analyses. ProteoMill overcomes this problem by checking in bi-weekly intervals if any external data sources have been updated, and if so, collects and processes the necessary external data.

Importantly, ProteoMill is for the most part based on *existing* R packages for statistical analysis and pathway annotation that are standard in the field. However, these methods are strongly focused on estimation of p-values and classifications of results based on p-value thresholds. This is an unfavorable approach to use of statistical methods and there is a need to move further in better estimation methods and expressing uncertainty.^19^ The methods for biological interpretation of complex pathways need preferably to be able to take into account the size of the effects and express the results communicating the uncertainty around the identification of pathways and their potential biological relevance. We plan that ProteoMill will be rapidly updated as such analytic methods and tools becomes available in the future.

In conclusion, the integrated features in this software provide powerful visualization strategies for the exploration of omics data, with a particular focus on the management and manipulation of proteomics data. By using this platform, researchers can expect to discover biologically relevant rendering of their data through results aggregated from reliable and up-to-date data sources.

The software offers innovative strategies to interactively explore quantitative proteomics data in a comprehensive workflow from data-upload to network analysis. It has a strong focus on well-maintained data sources, computational efficiency and user-friendliness, with in-app documentation that guides the researcher through the analysis.

## Methods

### Architecture and implementation

The software is developed in R (version 3.6.1) and the interface was created using the R-package Shiny and shinydashboard (version 0.7.1) with a customized CSS theme. Animations were created using jQuery and the library animejs.

### Identifier conversion

The Bioconductor package AnnotationDbi and EnsDb.Hsapiens.v86 was used to identify identifier type and for identifier conversion.

### Data quality control

2D PCA was implemented using the R-package factoextra and 3D PCA was implemented using the R-package Plotly. The software does not perform any assessment if the uploaded data fulfill the assumptions of these methods and it is the user’s responsibility to verify that the assumptions are fulfilled and interpret the results thereafter.

### Differential expression analysis

Two R-packages, limma^14^ and DESeq2^15^ were implemented for differential expression analysis. Each package is commonly used for fitting gene-wise linear models to expression data. limma was originally developed with a primary focus on the analysis of microarray data, while DESeq2 for the analysis of RNA-seq data and is based on the negative binomial distribution.

Differential expression analysis is conducted by specifying two contrasts and choosing a paired or non-paired design. The results are evaluated by inspecting the table in the “Differential expression” tab.

The results are displayed as estimated by the specific software, using the software’s default settings for shrinkage parameters, correction for multiple testing, significance level and etc.. For example, the correction for multiple testing is done using the Benjamini-Hochberg method and is applied to the tests performed within one run of the analysis and not with respect to all tests performed within one family of hypotheses in a study, which sometimes may be misleading.^20^ Further, the confidence intervals derived are not corrected for multiple testing. The user needs to verify if these setting are appropriate for the specific analysis done.

### Functional enrichment and network analysis

The hypergeometric distribution was used to calculate the probability of gene list overlap.

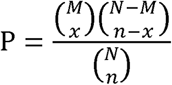

In this formula, N is the total number of genes in the background distribution, M is the number of genes in the background distribution annotated to a pathway, n is the total number of selected genes of interest, and x are the genes of interest annotated to a pathway.

Pathway data is dynamically collected from Reactome^21^ (https://reactome.org/download-data). MD5sum hashes are used to ensure that the local database is up to date.

Only lines where ‘Homo sapiens’ was listed in the ‘Species’ column, and ‘TAS’ (Traceable Author Statement) was listed in the ‘Evidence code’ column was kept.

For each entry in the main pathway data file, the top-level parent pathway was annotated. This was done by creating a directed acyclic graph (DAG) object using the R-package igraph^22^.

### Visualization of enrichment results

#### Volcano plot

In the volcano plot (Fig. 5b), each protein was annotated with the most top-level parent of the enriched pathway with the lowest adjusted p-value in which it occurs. Proteins labels were added for proteins with an absolute value of base-2 log fold change greater than 3.5. Label overlap was avoided using the geom_text_repel function in the ggrepel R-package.

A dashed horizontal line indicates which proteins had an adjusted p-value<0.05.

#### Sankey diagram

The differentially expressed proteins were visualized using a Sankey diagram (Fig. 6) in order to show in which over-represented pathways up- and down-regulated proteins occur in. The Sankey diagram was created using the R library networkD3, and was constructed to display three levels:

**Figure 6.**
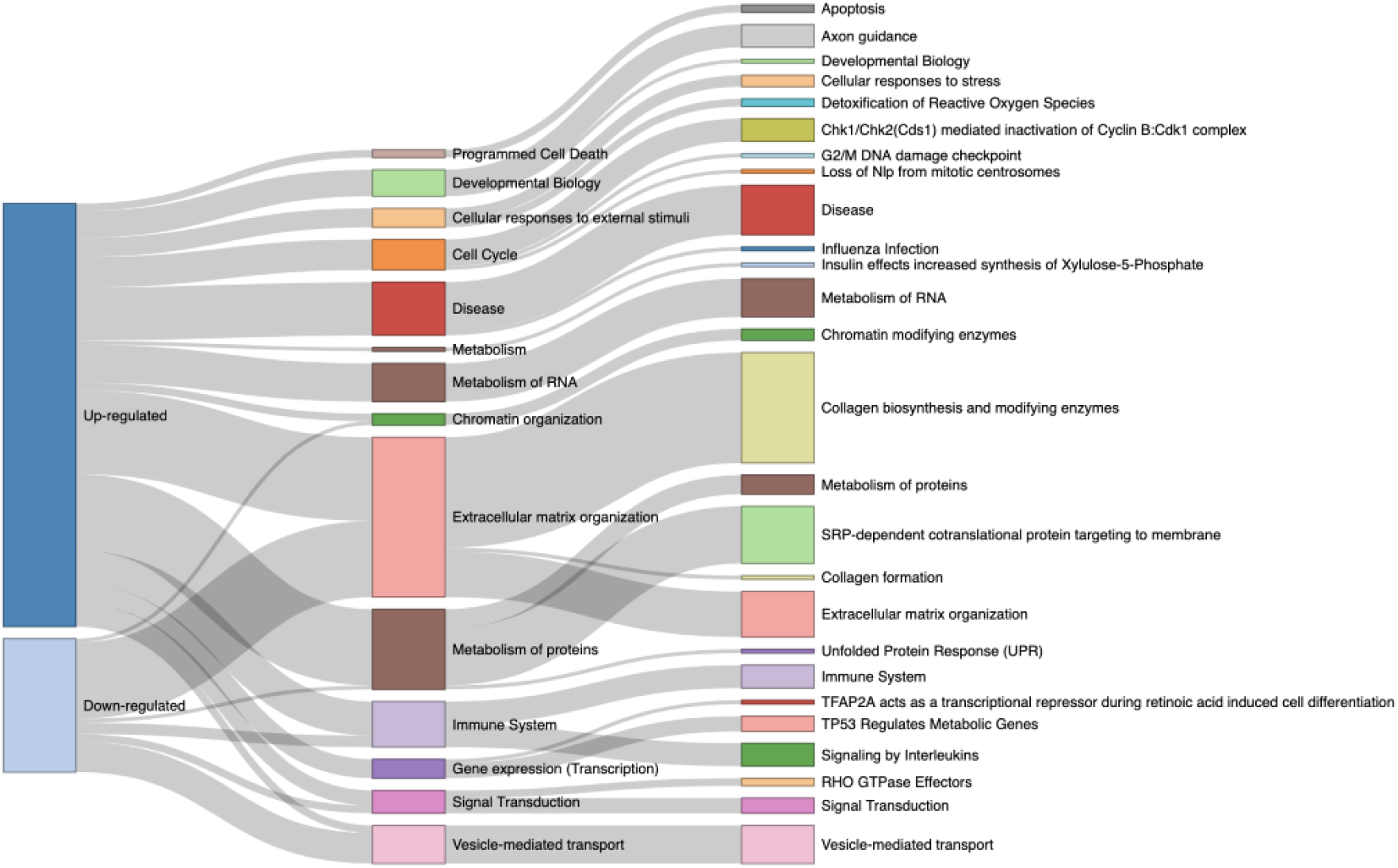
A Sankey diagram showing the distribution of up- and down-regulated proteins in highest and lower level Reactome pathways.

1. The division of up- and down-regulation of the proteins
2. The top level of the over-represented pathway
3. The pathway level that was over-represented.

### Network analysis

Protein-protein interaction (PPI) data was downloaded from STRING-DB^12^ (https://string-db.org/cgi/download.pl). Interactions are visualized using the visNetwork R-package. Node color is set according to the most significantly over-represented pathway to which a protein is annotated.

### Data sources

An important aspect of this software is to maintain data sources up to date. This is done by using an automated workflow at a bi-monthly interval. Data is collected from the two primary data sources, Reactome^21^ for pathway data and STRING^12^ for protein interaction data. These data are then structured to a predefined format, making it possible to integrate them in the analysis.

### Availability

The tool is freely available on https://proteomill.com/. Source code is available upon reasonable request.

## Acknowledgements

The authors would like to thank Aleksandra Turkiewicz, PhD for helpful discussions and critical reading of this paper.

